# Interactions between strains govern the eco-evolutionary dynamics of microbial communities

**DOI:** 10.1101/2021.01.04.425224

**Authors:** Akshit Goyal, Leonora S. Bittleston, Gabriel E. Leventhal, Lu Lu, Otto X. Cordero

## Abstract

Genomic data has revealed that genotypic variants of the same species, i.e., strains, coexist and are abundant in natural microbial communities. However, it is not clear if strains are ecologically equivalent, or if they exhibit distinct interactions and dynamics. Here, we address this problem by tracking 10 microbial communities from the pitcher plant *Sarracenia purpurea* in the laboratory for more than 300 generations. Using metagenomic sequencing, we reconstruct their dynamics over time and across scales, from distant phyla to closely related genotypes. We find that interactions between naturally occurring strains govern eco-evolutionary dynamics. Surprisingly, even fine-scale variants differing only by 100 base pairs can exhibit vastly different dynamics. We show that these differences may stem from ecological interactions in the communities, which are specific to strains, not species. Finally, by analyzing genomic differences between strains, we identify major functional hubs such as transporters, regulators, and carbohydrate-catabolizing enzymes, which might be the basis for strain-specific interactions. Our work shows that strains are the relevant level of diversity at which to study the long-term dynamics of microbiomes.

## Introduction

In nature, microbial communities contain individuals on a continuum of phylogenetic diversity, where both evolutionarily distant and proximate members coexist (*1, 2*). Members in the same communities can belong to different domains of life, such as archaea, bacteria, and fungi, and at the same time, can stably exist with extremely closely related relatives, a few single nucleotide polymorphisms apart (*3*–*6*). Collectively, closely and distantly related community members perform many vital ecological functions such as the regulation of biogeochemical cycles and fiber digestion in animal guts (*7, 8*). However, identifying how these different levels of diversity interact and co-evolve in complex communities remains an elusive and challenging problem.

A common strategy to study this problem is to analyze the dynamics of complex communities at different levels of diversity with metagenomic sequence data. Such data have been successfully leveraged to reconstruct the linkage between polymorphic sites in natural microbial populations, effectively allowing one to resolve both species and strain abundances (*9*–*12*). With these data, we can ask the important question: which level of diversity—strains, species (or even broader taxonomic units)—drives community dynamics, and through which factors—those intrinsic to the community or extrinsic from it? Differentiating between dynamics induced by intrinsic factors, such as biological interactions, and those induced by extrinsic factors, such as environmental changes, requires careful control of the environment, which is rarely possible in natural systems such as mammalian guts. Such a level of control can however be achieved *in vitro*, by domesticating natural communities in controlled laboratory conditions (*13*). Domesticating natural communities also allows us to study multiple community replicates in the same abiotic environment, thus helping us disentangle which outcomes of community dynamics are repeatable and general, and which ones are chance and context specific.

Here, we show that interactions between strains—which are genetic variants of the same species— govern long-term microbial community dynamics. These results stem from tracking ten domesticated microbial communities from the pitcher plant *Sarracenia purpurea* for more than 300 generations. By analyzing dynamics at several phylogenetic levels, we show that community composition varies most strongly at the level of strains. Further, interactions within each community were strain-specific: at times decoupling the dynamics of one strain from its close relative. Remarkably, as few as 100 base-pair differences across strain genomes (∼99.99% similarity) were sufficient for strain dynamics to diverge from each other. Finally, we show that strains can differentiate by fine-tuning only a handful of functional categories in their genomes, such as transcriptional regulators, metabolite transporters and tricarboxylic acid (TCA) cycle enzymes. Together, our results highlight that strains may be the relevant unit of interaction and dynamics in microbiomes, not merely a descriptive detail.

## Results

### Naturally occurring strains coexist for hundreds of generations in laboratory microcosms

We followed the eco-evolutionary dynamics of 10 replicate microbial communities derived from the carnivorous pitcher plant *Sarracenia purpurea* (Fig. 1a). We began by sampling communities from 10 distinct pitchers from plants belonging to the same bog, and after sampling, filtered out particles larger than 3 μm (to focus on bacteria). We then transferred and propagated each replicate community through serial passaging in a medium consisting solely of acidified water and ground cricket powder as the nutrient source (see Methods). This medium mimics, in part, the ecological conditions of the pitchers from which we derived these microbes.

**Figure 1:**
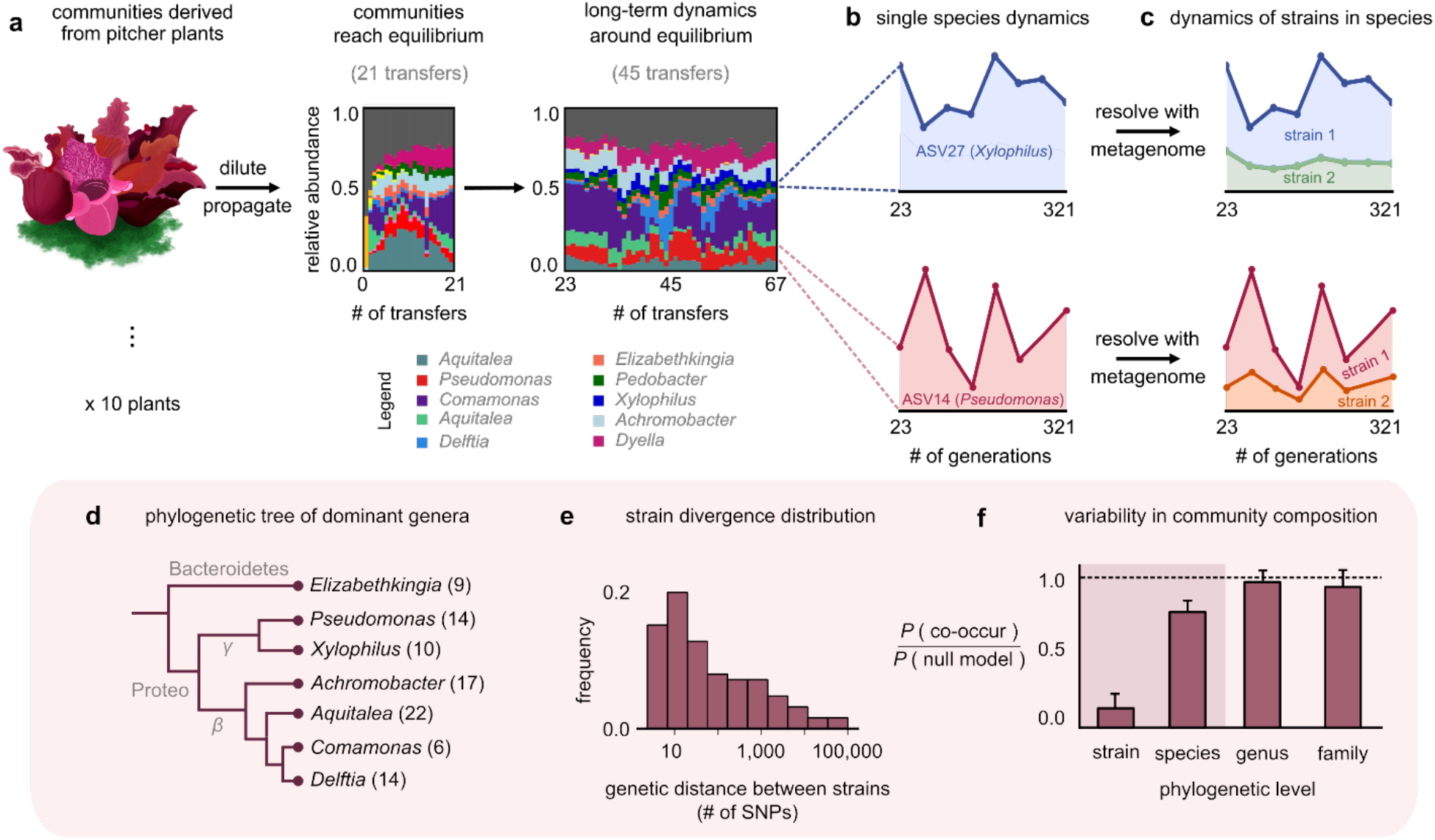
Closely related strains coexist for hundreds of generations in pitcher plant-derived microbial communities. **(a)** Diagram illustrating our experimental protocol. Stacked bar plots show the composition of 1 community (M06) at the ASV (species) level sampled at each transfer; each color corresponds to a unique ASV which we tracked further using metagenomic sequencing, with their genera in the legend. [Illustration credit: Michelle Oraa Ali.] **(b)** Relative abundances of ASV27 (blue) and ASV14 (pink) with the approximate number of generations (see Methods); the 8 time points shown correspond to those for which we collected metagenomic data. **(c)** Relative abundances of strains identified in ASV27 and ASV14 using metagenomes. The shaded colors correspond to the abundance of each of the two strains. **(d)** Phylogenetic tree of the dominant strain-containing taxa across all 10 communities, with the identified genus names labeled. Brackets indicate the number of detected strains belonging to each genus. **(e)** Distribution of the genetic distance (divergence) between strains belonging to the same species, measured in the number of detected single nucleotide polymorphisms (SNPs) differentiating them. **(f)** Bar plot showing the probability with which two members of the same taxonomic group (family, genus, species, etc.) co-occurred in a sample, normalized with a null model where all members were distributed randomly across communities. Dashed line indicates the null expectation.

To stably maintain high diversity, we initially propagated communities at a low dilution factor (1:2) every 3 days for 21 transfers, during which they reached distinct, rich and stable equilibria (Fig. 1a) (ecological dynamics studied in a previous publication (*14*)). To study community dynamics over evolutionary timescales, we followed them using a similar protocol, but at a higher dilution factor (1:100) and less frequent transfer rate (every 7 days) for more than 300 generations (45 additional transfers). To resolve both coarse (species) and fine (genotype) scale dynamics, we performed 16S rRNA and deep metagenomic sequencing, respectively. Specifically, we used 16S rRNA sequencing to follow species composition (with species defined as having identical 16S rRNA); this gave us the “ecological” dynamics of each community. Metagenomic sequencing, which we performed at 8 evenly spaced time points between transfers 23 and 67, allowed us to follow “evolutionary”, or genotype-level, changes in each community (Fig. 1c). Together, this setup enabled us to track the eco-evolutionary dynamics of all 10 communities.

At the species level (variants with the same 16S rRNA sequence), the composition of each community fluctuated dynamically, but remained around the same equilibrium state for >300 generations (Fig. 1a; median ∼31% temporal coefficient of variation in the abundance of a species; Fig. S1). We hypothesized that genetic variation occurring during the experiment, through mutations and recombination, might be responsible for these fluctuations. To test this, we mapped metagenomic reads from each sample to a database of 33 reference genomes belonging to isolates from our communities (see Methods). To avoid sequencing-related artifacts, we ensured that read mapping was competitive, i.e., we only used those sequenced reads that mapped unambiguously to one genome. Importantly, because our reference genomes belonged to isolates from our communities, any detected genetic variation indicated the presence of variants relative to a resident of these communities.

Most (97%) genetic changes were in the form of single nucleotide polymorphisms (SNPs), an overwhelming majority (∼98%) of which were detected in the first sequenced time point (∼23 generations; Fig. S2). This suggested that the communities had significant pre-existing, or standing, genetic variation. Given the large degree of divergence (∼1% genome-wide divergence for some pairs; Fig. 1e), the pre-existing variation likely came from genetic variants in the plants, rather than from variants arising during the first 23 generations of lab propagation. Therefore, we rejected our original hypothesis that mutations and recombination occurring during the experiment were the main driver of community dynamics. Instead, we focused on the dynamics of pre-existing variants. Almost all (∼99%) SNPs were bi-allelic (had only 2 variants; Fig. S2), which allowed us to track variant dynamics in terms of the temporal trajectory of each SNP.

The temporal trajectories of SNPs in the same reference genome (species) were highly correlated, with their allele frequencies increasing and decreasing together (mean correlation coefficient 0.8; *P* < 0.01; Fig. S3). Such highly correlated allele frequency trajectories are hallmarks of genetic linkage (*9, 15*), and suggest that the large number of SNPs in each species were co-localized in a small number of genotypes or strains. We clustered the allele frequency trajectories and could statistically detect at most 2 strains for each species (see Methods and Fig. S4). For each cluster (strain), we estimated its frequency within the species as the mean allele frequency of the cluster (see Methods). We calculated the relative abundance of each strain by multiplying its frequency with the relative abundance of the species it belonged to. Together, we concluded that within each community, there was a pair of strains underlying most taxa, including the phylogenetically diverse *Elizabethkingia, Aquitalea*, and *Delftia* (Fig. 1d). Within each species, strain dynamics displayed vastly different patterns. For some species, only one of the strains changed appreciably in abundance over the experiment (Fig. 1c, top; *Xylophilus sp*.). For other species, both strains fluctuated constantly throughout the experiment (Fig. 1c, bottom; *Comamonas sp*.).

Since we observed different conspecific strains in different communities, we next asked at which phylogenetic level the 10 replicate communities were most variable at, i.e., strains, species, genera, or families. For this, we calculated the probability that two members of the same taxonomic group (family, genus, species, etc.) co-occurred in a sample, normalized against a null model where all members were distributed randomly across communities (see Methods). We found that community composition across all 10 replicates was much more variable at the level of strains than species (Fig. 1f). This result is consistent with previous work that natural microbial communities are taxonomically variable at finer phylogenetic levels (strains and species), but more similar at coarser levels (families) (*16, 17*). Together, we concluded that natural strains of the same species can coexist in microbial communities over hundreds of generations, maintaining the communities in distinct stable states.

### Highly related strains can decouple in their dynamics

Motivated by the observation that closely related strains persisted in communities, we asked whether the dynamics of each strain were similar, or coupled, to that of their closest relatives. To answer this question, we measured a strain-strain coupling for strains of the same species, defined as the correlation between their abundance trajectories (Fig. 2a and Methods). Strain dynamics could either be highly correlated (“coupled strains”, Fig. 2a) or be uncorrelated (“decoupled strains”). Measuring strain-strain coupling for each species across all 10 communities revealed a bimodal distribution, whose two modes were occupied by highly correlated and uncorrelated strain pairs, respectively (Fig. 2b). The shape of this distribution allowed us to reliably classify strain pairs as either decoupled (coupling < 0.4, the inflection point of the distribution; see Methods) or coupled (coupling > 0.4). We thus concluded that ∼20% of conspecific strains, belonging to the same species, displayed rather dissimilar or decoupled dynamics.

**Figure 2:**
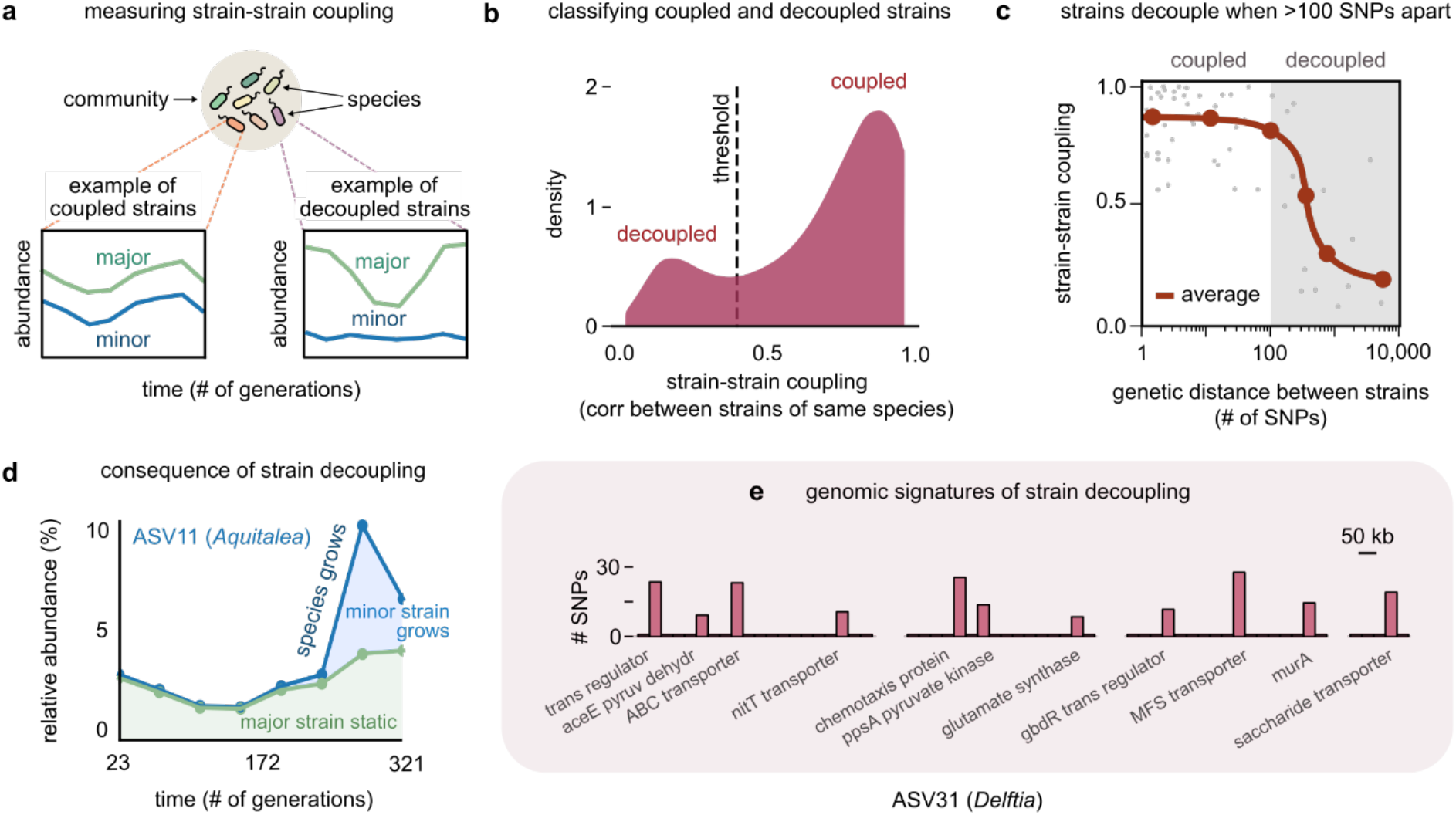
Even highly related strains (∼100 SNPs apart) can decouple in their dynamics. **(a)** Schematic showing examples of strain-strain coupling. We defined strain-strain coupling as the temporal correlation between strain abundances belonging to the same species in a community; coupled strains (left) had highly correlated abundances, while decoupled strains (right) were uncorrelated. **(b)** Distribution of the strain-strain coupling across all species and communities; dashed line shows the threshold coupling used to classify strains as coupled and decoupled (see Methods). **(c)** Strain-strain coupling as a function of the genetic distance between strains. Each gray point represents a conspecific strain pair. The solid red line shows a moving average (LOESS fit). **(d)** Relative abundance of ASV11 of the genus *Aquitalea* in community M04, and its underlying strains over time; the blue shaded region represents the minor strain, while the green region represents the major strain. The solid blue line shows the total species abundance. **(e)** Bar plot showing the number and genetic location of SNPs in the core genome of ASV31 from community M04, whose strains were decoupled and differed by 186 SNPs. Each bar shows the number of SNPs in one gene along with its annotation. Only SNPs belonging to annotated genes are shown.

To test if the genetic distance between strains influenced their strain-strain coupling, we plotted the average coupling as a function of the genetic distance between a conspecific strain pair (see Methods). Remarkably, the average strain-strain coupling decreased sharply beyond a genetic distance of just about 100 SNPs (Fig. 2c). This suggested that changes in as few as 100 base pairs, corresponding to roughly 0.01% of these genomes, were sufficient to decouple strain dynamics. A consequence of such decoupling was that in species with decoupled strains, drastic changes in the species’ abundance were driven only by changes in one of the strains (Fig. 2d and Fig. 1c, top).

To understand the genomic signatures of strain decoupling, we examined the location and putative function of SNPs in a pair of highly related but decoupled strains (see Methods). The *Delftia* genome in community M04, which had the fewest (186) SNPs among its decoupled strains, shed light on which genes were associated with decoupling. The SNPs differentiating these strains were scattered throughout the core genome, with 62% in the coding regions of genes with known functional annotations. Broadly, these genes corresponded to transcriptional regulators such as *gbdR*, transmembrane proteins such as the nitrate transporter *nitT*, and enzymes implicated in central carbon metabolism, such as *aceE, murA* and *ppsA* (Fig. 2e). The *gbdR* protein is a known transcriptional regulator of amino acid metabolism, glycine betaine catabolism as well as phosphatase activity in bacteria, and may play a role in differentiating metabolic flux balance between strains (*18*). The *nitT* protein is member of the major facilitator superfamily (MFS), and a transporter that controls the uptake of nitrate available in the environment (*19*). Finally, *ppsA* and *murA* are enzymes that metabolize pyruvate and its derivatives, and may determine how strains catabolize metabolic intermediates before they enter the TCA cycle (*20, 21*). Together, this evidence suggests that strains can decouple even by diversifying in only a handful of functional pathways.

### Community interactions are strain-specific

Equipped with the understanding that conspecific strains could display decoupled dynamics, we asked which ecological factors were responsible for the observed differences in strain dynamics. Our experimental setup explicitly controlled for abiotic ecological factors, since all communities were grown in the same environmental conditions. Therefore, any dynamical differences between strains must have emerged as a result of biotic factors, such as ecological interactions. To detect and measure interactions between community members, we exploited the fact that all 10 communities were at equilibrium (Fig. 1a). In a controlled environment, interactions between community members at equilibrium are expected to induce temporal correlations between their abundances (*22*–*24*). In contrast, we expect almost no correlations if all abundance fluctuations are purely stochastic (due to neutral drift) (*25*–*28*). Motivated by this logic, we used temporal correlations between the abundance trajectories of pairs of community members to detect the presence and strength of possible interactions between them.

We first looked for the presence of interactions between members for each community separately. To do this, we inferred two interaction networks: one at the level of species and the other at the level of strains. To infer an interaction network, say at the species level, we measured all pairwise correlations between the abundances of species in the community. Two species were said to interact if the correlation between them was statistically significant compared with an ecological neutral model. Briefly, this model computed the expected distribution of correlations between non-interacting members of a community by simulating their abundance trajectories under known empirical laws, using only the mean and variance of their observed relative abundances (see Methods). Importantly, even at the level of strains, we specifically looked for interactions between strains belonging to different species, not between strains of the same species. To illustrate our results, we focused on community M07 as an example (Fig. 3a). We found that the strain-level interaction network of this community was 90% denser in interactions (38 interactions across 24 strains; density measured as number of interactions per node) than the species-level network (10 interactions across 12 species). In fact, this observation of denser strain-level interaction networks was true for all communities (mean ∼70% denser networks at the strain-level). Most strain-level networks revealed interactions which could only be observed at the level of strains, not species (e.g., Fig. 3a, *Rh*). This suggested that in terms of the presence/absence of interactions, community interactions were likely strain specific.

**Figure 3:**
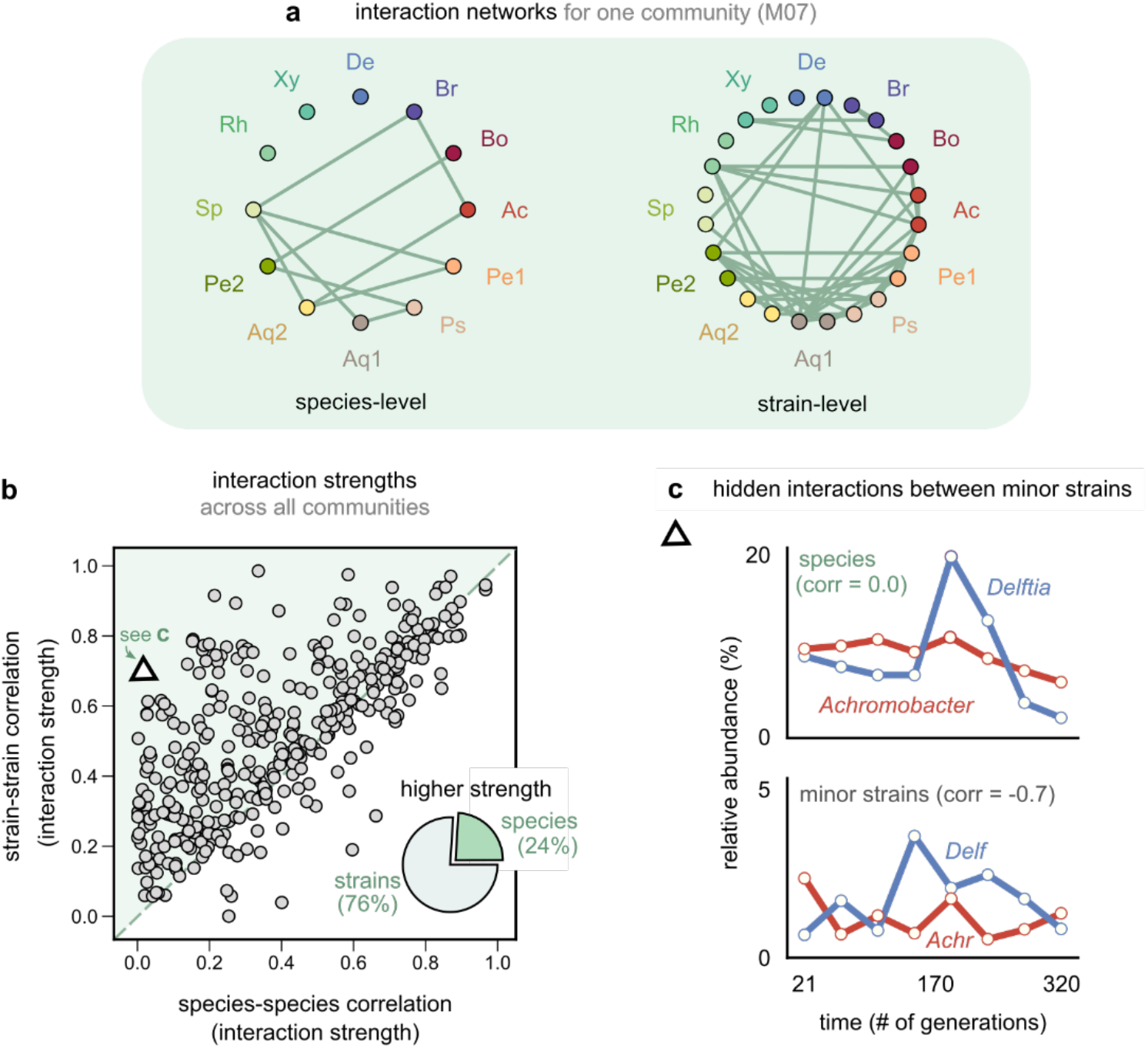
Community interactions are strain-specific. **(a)** Interaction networks inferred using dynamical correlations from community M07, measured at the species (left) and strain level (right) (see Methods). Each node represents a species (left) or strain (right), and link indicates the presence of a detected interaction; Ac: *Achromobacter*, Pe: *Pedobacter*, Ps: *Pseudomonas*, Aq: *Aquitalea*, Sp: *Sphingomonas*, Rh: *Rhodobacter*, Xy: *Xylophilus*, De: *Delftia*, Br: *Brevundimonas*, Bo: *Bosea*. **(b)** Scatter plot of the dynamical correlation between species in a community and the highest correlation between their corresponding strain pairs. Each point represents one species in one of the 10 communities. The shaded region indicates strain-strain correlations higher than species-species correlations. Inset shows a pie chart of the fraction of points supporting higher strain-level interactions (76%) versus species-level interactions (24%). Triangle indicates a pair of *Achromobacter* (red) and *Delftia* (blue) species, shown in **(c)** [top] Relative abundance plots of two uncorrelated species measured over the experiment; [bottom] Relative abundances of the minor strains for the same species, which are strongly negatively correlated.

Having looked at the presence of interactions, we next focused on interaction strengths. To do so, we measured the magnitude of the correlation coefficients between taxa in the same community as proxies for their interaction strengths. Remarkably, we found that for most taxa (76%), at least one pair of strains, belonging to two different species, was more strongly correlated than the corresponding species themselves (Fig. 3b; *P* < 10^−6^, Wilcoxon-signed rank test adjusted for multiple comparisons). To test if this result could arise purely because we compared four pairs of strains for each pair of species, we used a null model in which strains were randomly sampled and combined into new mock “species”. (see Methods). This shuffling coalesced unrelated pairs of strains, but preserved the number of comparisons performed, and revealed that the observed high fraction of strain-dominant interactions was not a statistical artifact (*P* < 10^−3^, permutation test; Fig. S6). Therefore, not only were putative strain-level interactions more numerous in communities (Fig. 3a), when present, they were also much stronger (i.e., even when both strain and species level correlations were high, the values of at least one pair of strain correlations were often higher; Fig. 3b). Finally, in 7% of cases, our analysis revealed so-called “hidden” interactions, masked at the species level but visible with strains. In these cases, while there was virtually no correlation between species dynamics (Fig. 3c; Pearson correlation 0.02, *P* = 0.9), the dynamics of their minor strains were strongly correlated (Fig. 3c; Pearson correlation -0.71, *P* < 0.01). Together, these data suggest a striking tendency for community dynamics to be driven by ecological interactions between strains, not species.

### Genetic variation in transporters, regulators and pseudogenes differentiate strains

Having observed that strains, not species, govern the eco-evolutionary dynamics of communities, we finally asked what differentiated strains at the genomic level. We had already explored a specific example while studying decoupled strains (Fig. 2e); we now asked for more general features that differentiated coexisting strains. Most (97%) of the differences between strains in their core genomes were in the form of SNPs; the remaining 3% constituted insertions and deletions (average 4 bp per event; see Methods). Since we had only one reference genome for each ASV, we could not detect changes in the flexible genome, such as gene gains in one strain compared to the other. For all 163 strains, there was no significant bias in the distribution of SNPs between coding and non-coding regions, implying that most (>80%) of the SNPs were found in coding regions (Fig. S7).

We next asked which functional categories of genes were enriched in strain-differentiating SNPs. For this, we performed a broad categorical analysis, giving us a bird’s eye view of the cellular functions likely to be variable in strains (we used annotations and categories from the KEGG database, respectively; see Methods) (*29*). Four major functional categories differentiating strains emerged: (1) two-component systems, (2) enzymes involved in carbon metabolism, (3) transcription factors and (4) transporters (Fig. 4a). To get a closer look at specific examples of pathways affected by each of these four categories, we examined ASV11, belonging to the genus *Aquitalea* in M04, which contained SNPs in genes from all four categories (Fig. 4b; Table S2). For two-component systems, we found the *dctD* protein, which along with *dctB*, regulates the uptake of C4-dicarboxylates such as aspartate, malate, fumarate and succinate (*30*). The enzyme *korB* is an oxidoreductase, known to decarboxylate 2-oxoglutarate, a key intermediate in the TCA cycle (*31*). The transcription factor *phnF* represses the *phnCDE* operon and regulates the biosynthesis of amino acids such as glutamate (*32*). Finally, we found many diverse sugar and amino acid transporters (of the ABC and MFS families), ion transporters like *corA* (*33*), as well as *iutA*, which mediates siderophore uptake and competition for iron in bacteria (Fig. 4b) (*34*). These examples typify some key functional signatures that differentiate strains, and suggest that variation in transporters, regulators and enzymes in central carbon metabolism may be the basis for strain-specific ecological interactions, e.g., by separating the metabolic niches of strains.

**Figure 4:**
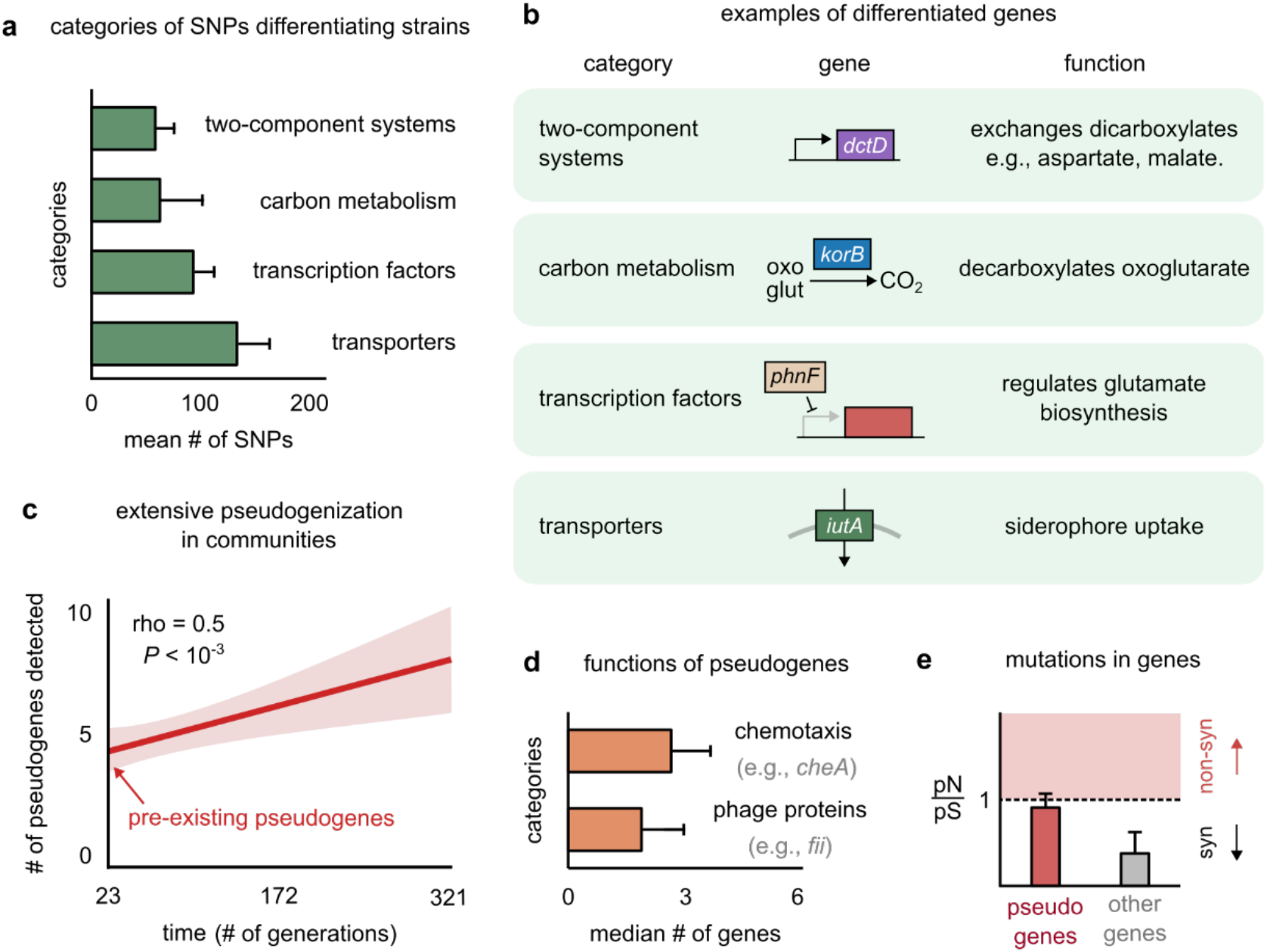
Genetic variation in regulators, transporters and pseudogenes differentiate strains. **(a)** Bar plot showing the four functional categories of genes most enriched in strain-differentiating SNPs. The x-axis represents the mean number of SNPs belonging to the category in a strain pair. **(b)** Table showing an example of a gene in each functional category identified in (a); the middle column shows a schematic of the gene with its name (italics). **(c)** The average number of pseudogenes detected in strains per strain as a function of time. The solid line shows a linear regression, whose intercept shows the number of pseudogenes detected at the first sequenced time point; the shaded region represents the s.e.m. **(d)** Bar plot showing the two functional categories most enriched in strain-differentiating pseudogenes. The x-axis represents the median number of genes belonging to the category in a strain pair. **(e)** Bar plot showing the mean pN/pS of mutations detected in pseudogenes (red) and all other strain-differentiating genes (gray). Dashed line represents the expected pN/pS under a neutral model. All error bars represent s.e.m.

Another key genomic signature was the presence of several strain-specific pseudogenes in strain genomes. Pseudogenes are genes that contain a premature stop codon and are not expected to produce functional proteins (*35*). Not only did strains have several (∼5) pseudogenes at the first time point (∼23 generations), but we could detect new pseudogenes throughout the experiment (Fig. 4c; see Methods for how we detected pseudogenes). This suggested that pseudogenization was both extensive and rampant during community evolution. Pseudogenes were enriched in two functional categories: (1) motility proteins such as *cheA*, responsible for chemotaxis (*36*), and (2) phage proteins such as *fii* and *gpdD*, known to constitute viral tails (Fig. 4d) (*37*). Functions such as chemotaxis and structural viral proteins are not expected to be advantageous for bacterial growth in the environmental conditions of our experiment, and are thus likely to face weak, or no selection. Indeed, while pseudogenes did not accumulate mutations faster than other genes (Fig. S8), their mutational profile had a pN/pS ∼ 1, consistent with neutral evolution (Fig. 4e). In contrast, other genes showed signatures of purifying selection, with a pN/pS ∼ 0.2, significantly lower than 1 (Fig. 4e). The rampant pseudogenization observed in our communities suggests that evolution tends to deactivate genes that do not contribute to fitness, such as chemotaxis and viral genes in stable, well-mixed environments. The end products of such non-functional pseudogenization and functional genomic variation may be strain-specific interactions, allowing even highly related strains to show decoupled dynamics.

## Discussion

Understanding the role of strains in microbiomes is crucial to the study of microbial ecology and evolution (*38*). Here, we addressed this question by propagating microbial communities from pitcher plants for more than 300 generations. We found that strains, which were pre-existing genetic variants belonging to the same species, were the key determinants of long-term community dynamics. Differing by between 1 and ∼10,000 SNPs, strains imparted both compositional and dynamical variability to communities. Compositionally, we found that communities were most variable in terms of strains, with each community carrying a unique set of strains even when they carried the same species. This variability was likely the result of two chance events: stochastic sampling and laboratory selection. Dynamically, once the communities settled into unique equilibria, we found that the dynamics of even extremely closely related strains, with as few as 100 SNPs, could be decoupled from each other. Even when a species’ abundance appeared relatively stable over time, strains comprising it could be highly dynamic. Such strains were temporally correlated with other strains belonging to different species. This suggested that most putative ecological interactions in the communities were specific to strains, not species.

Our observation that interactions among existing strains govern evolutionary dynamics is perhaps somewhat different from one’s naïve expectation, anticipating new mutations and recombination events to be the dominant forces driving evolution within communities (*11, 39*). Once our communities reached a stable equilibrium, their evolution was marked by the dynamics of the pre-existing strains, which could stably coexist for hundreds of generations. Thus, in both cases, either by mutation and recombination (in single-species laboratory evolution) or by assembly (in complex communities as in this paper), the dynamics of populations are determined by the ecological interactions between fine-scale genetic variants, or strains (*9, 15*). Moreover, in both cases, the interactions between strains are likely resource-mediated, since the genetic differences between ecologically divergent strains are concentrated in metabolic genes (*11*). Taken together, these results suggest a striking parallel between the eco-evolutionary dynamics of both isogenic populations as well as complex communities.

The patterns we observed, such as strain decoupling and strain-specific interactions, underscore the emerging view that strains are the most dynamic and interactive units of microbiomes (*15, 40, 41*). Our study has one major advantage over previous work where natural communities, such as human gut microbiomes, were sampled over time without environmental control. That is that by experimentally propagating natural samples, we shielded the communities from external host-induced perturbations such as the migration of new genetic variants or immune system control. Therefore, any observed community dynamics were a result of intrinsic causes, such as strain-specific interactions, not extrinsic causes, such as host-induced shifts. The reproducibility of these patterns across 10 independent replicate communities further strengthens our findings.

Finally, our results raise a question about the appropriate level at which to monitor microbial community composition. On one hand, our results show that strains as few as 100 SNPs apart can have independent ecological dynamics and interactions, and as such, represent distinct ecological variables. On the other hand, consistent with previous work, our results also show that the presence of strains is highly variable across communities, even in the same abiotic environmental conditions (*40*). Therefore, coarse-grained assemblages (e.g., members of the same taxonomic family) should better represent the relationship between metabolic niches and community composition (*16, 17*). But it is not clear how such assemblages should be defined in general, and what type of functional redundancy they capture. This is because there are numerous definitions of what constitutes “function”. In the study that preceded this paper, we found that the substrate consumption profile of the community was strongly correlated with community composition (at the 16S rRNA level), indicating thus that there is, at best, weak functional redundancy in the community if substrate preference is what we define as “function” (*14*). However, if what we care about is the conversion of organic carbon to CO_2_, for instance, community composition is indeed highly redundant (most species in the community could perform aerobic respiration and yield similar amounts of CO_2_). More work is needed in this arena to clarify how phylogenetic resolution maps to relevant ecological functions. However, at the very least, we can say that if our questions pertain (eco)evolution, strains are the appropriate variables with which to study the system.

## Methods

### Experimental methods

#### Sampling, experimental design, and DNA extraction

We collected aquatic samples from 10 healthy *Sarracenia purpurea* pitchers at Harvard Pond (Petersham, MA), filtered them through 3 µm syringe filters, combined them in a 1:1 ratio with sterilized cricket media, and grew them in 48-well plates in a 25°C incubator. The sampling and experimental design was identical to, and is described in detail in, Bittleston et al (*14*). For the first 63 days (21 transfers), every 3 days each sample was mixed well and 500 µL was transferred to a new plate with 500 µL of sterile cricket media. After this point, we shifted to sampling every 7 days using a larger dilution ratio of 1:100; with 20 µL of each community mixed into 1,980 µL of cricket media. At every transfer, we froze a portion of each sample at −80 °C for later DNA extraction and sequencing. We extracted and quantified DNA as described in Bittleston et al. 2020 using the Agencourt DNAdvance kit (Beckman Coulter) the Quant-iT PicoGreen dsDNA Assay kit (Invitrogen), respectively.

#### Strain isolation

Individual strains were isolated from five of the ten microcosms (M03, M05, M07, M09 and M10) by plating the culture fluid and picking around 100 colonies per microcosm. Details of the isolation methods and preliminary strain identification are described in Bittleston et al (*14*). We chose a set of 33 diverse strains that were well-represented in the amplicon sequencing data from the first 63 days, and extracted DNA using the same procedure as the community samples (Supplementary Table S1).

#### Sequencing

Amplicons were sequenced at the Environmental Sample Preparation and Sequencing Facility at Argonne National Laboratory on a MiSeq targeting the V4 region of 16S rRNA using the 515F and 806R primers (*42, 43*), using the same procedure as described in Bittleston et al (*14*). Metagenomes from all 10 communities were sequenced across 8 evenly spaced time points, resulting in 80 samples. Genomic and metagenomic libraries were prepared at the BioMicro Center at MIT using NexteraFlex and sequenced on a NovaSeq SP500 run, aiming for 20x more sequencing depth for strains relative to metagenomes.

### Bioinformatic methods

#### Species abundance estimation

We defined each species in our analysis as a unique Amplicon Sequence Variant, or ASV. To analyze the amplicon sequencing data, we used the same ASV assignments and sequence analysis procedure described in Bittleston et al (*14*). Each species (or ASV) was identified with a unique ∼250 base pair long sequence. In each sample, we estimated the relative abundance of a species as the number of reads which mapped to it, normalized by the total number of reads in the sample. We estimated the number of elapsed generations after each cycle in the experiment as log_2_(*D*), with the dilution factor *D* = 100. Unless stated otherwise, we performed all subsequent analyses using Python v3.7.3 and the NumPy and SciPy packages (*44, 45*).

#### Reference genome database construction

We built a reference genome database by sequencing 33 out of the 100 isolates extracted from our communities (described above). After trimming raw sequencing reads with Trimmomatic 0.36 (*46*), we assembled genomes using SPAdes v3.13 (*47*). We removed all assembled contigs less than 200 bp in length. We then used the Genome Taxonomy Database (GTDB-Tk) to assign genus-level identity to each genome and to generate a phylogenetic tree (*48*).

#### Read mapping

After trimming reads with Trimmomatic 0.36, we mapped each read against our reference genome database using Minimap2 v2.17 (*49*). Since our reference genomes were assembled using isolates from the same communities as the reads, we used stringent settings for short reads (`-ax sr`) and to ensure unambiguous read mapping, only kept mapped reads with the highest MAPQ quality score of 60. Further, we filtered out any remaining reads which mapped equally well to more than one location in the reference database. We mapped reads sample by sample, each sample corresponding to a unique time point from among 1 of our 10 communities.

#### Variant calling

To identify genetic variants using the aligned reads, we used the Bayesian genetic variant detector FreeBayes v1.3.2, which prioritizes variation in literal read sequences over variation in their exact alignment (*50*). To calibrate the variant caller for haploid prokaryotic genomes, we used the settings `--ploidy 1 --haplotype-length 0 --min-alternate-count 1 --min-alternate-fraction 0 --pooled-continuous --report-monomorphic -m 60 -e 1000`. For downstream analysis, we only considered bi-allelic (only one alternate allele observed at the site, >97%) SNPs which had a Phred quality score ≥ 20, and a local read depth ≥ 10 (i.e., allele frequency least count: 10%). To avoid sequencing errors and read mapping artifacts, once a SNP was detected at a certain time point in a community, we required that it be detected across all subsequent time points, or until the time point at which the species containing it reached a relative abundance of zero.

#### Strain identification

For each community, we chose all SNPs detected at the first time point and partitioned them among the 33 reference genomes (unique 16S rRNA sequences, or species) they were detected in. For each set of SNPs detected in a species, we clustered the frequency trajectories of their major allele (reference or alternate allele, whichever was higher) using dynamic time warping, a generalized *k*-means clustering algorithm for time-series data (*51*). We performed clustering for 1, 2 and 3 clusters for each species. In all cases, a third cluster had low Euclidean distance (mean < 0.15) from and was visually indistinguishable from one of the other two clusters (Fig. S4). We thus rejected it. A second cluster either did the same or comprised a set of SNPs which with nearly fixed allele frequencies (> 0.9). We inferred the latter clusters as belonging to those loci that were shared by all conspecific strains in the sample but differentiated them from the reference genome. If the latter type of cluster was present, we assigned 2 clusters to each species. If not, we assigned 1 cluster. The first 2 clusters always accounted for > 80% of SNPs in a species, which increased our confidence in cluster assignment. Clustering using an alternate method, such as UPGMA (unweighted pair group method with arithmetic mean) clustering on the data (using the Euclidean distance between SNP trajectory correlations as a distance metric), yielded similar results (Fig. S5). Always, one cluster consisted of SNPs differentiating the two strains, and the second cluster consisted of shared SNPs between them.

#### Strain abundance estimation

We identified each strain as being represented by its reference genome along with the cumulative set of SNPs in both clusters. To assign each strain as major or minor, we estimated their frequency within each species. For this, we only used the allele frequencies of SNPs in the first cluster. Specifically, we calculated the average allele frequency weighted by the local read depth for each SNP in the cluster. The strain with a higher weighted mean frequency at the first time point was classified as the major strain. The abundance of each strain was calculated as its frequency at the time point (between 0 and 100%) times the relative abundance of the species in the community at that time point. Thus, each strain pair partitioned the abundance of its species into two abundance trajectories, that of a major and a minor strain.

#### Functional annotation of strain-differentiating SNPs

We performed a categorical enrichment analysis of strain-differentiating SNPs (first SNP cluster, described above). First, we extracted those SNPs detected in the genes of each reference genome, on average 82% of SNPs. We annotated the proteins corresponding to each gene using eggNOG-mapper v2 (*52*), with parameters `--go_evidence non-electronic --`**target_orthologs all -- seed_ortholog_evalue 0**.**001 --seed_ortholog_score 60**. After filtering out genes with unknown functional annotations such as hypothetical proteins and proteins with domains of unknown functions, as well as genes with no known KEGG Orthology (KO) group (30% of SNP-containing genes), we asked which functional categories of genes had more SNPs than expected by chance. Each KO was associated with a specific functional category in the associated BRITE hierarchy. For the BRITE category “Enzymes”, which is extremely broad, we manually chose finer functional categories, such as carbon, nitrogen and sulfur metabolism, based on what compounds the enzymes acted on. For each functional category containing at least 1 SNP, we calculated an enrichment score, given by the inverse of the *P* value of the observed number of SNPs in the category when compared against the expected number of SNPs (given by a binomial distribution with success rate equal to the fraction of the genome corresponding to that category of genes). Gene annotations and names in examples were derived manually using the KEGG and UniProt entries corresponding to each gene’s KO numbers.

#### Pseudogene detection and analysis

To detect pseudogenes, we used those strain-differentiating SNPs which were localized in genes (described above). Specifically, we identified the codon-level changes engendered by each SNP, and a pseudogene was identified as one which resulted in a premature stop codon. Like all SNPs, we required that this stop codon-enabling SNP be detected at all subsequent time points once detected. For functional analysis, we used the same procedure as described above, but restricted to pseudogenes. To detect mutations in genes, we counted those SNPs which were not detected at the first metagenomic time point but detected at later time points; like other SNPs, once detected, we required mutations to be repeatably detected at all subsequent time points. To calculate pN/pS for each gene (at the final time point), we accumulated all SNPs detected in that gene and classified them as synonymous or non-synonymous based on whether they led to an expected amino acid change; we then calculated pN/pS using standard techniques.

### Statistical methods and models

#### Community compositional variability

We measured the variability in community composition across our 10 communities at different taxonomic levels. At each taxonomic level, we partitioned all community members into groups; members within a group differed at that taxonomic level but shared a common ancestor just one taxonomic level above. As an example, to measure variability at the species level, we partitioned all members into groups; each group contained different species belonging to the same genus (say all 3 species belonging to the genus *Aquitalea*). After partitioning, we measured the probability with (or frequency at) which two members of a group co-occurred in a sample (how often two different *Aquitalea* species co-occurred). This gave us a measure of how different two communities were at a given taxonomic level (in this example, to what extent different communities had different species of the same genus).

To normalize the observed probability of co-occurrence against the expected probability at each level, we repeated the calculation by randomly shuffling member labels across groups. This procedure destroyed any phylogenetic relationship between members of a group, but preserved both the number of groups and their sizes.

#### Strain-strain coupling

We measured a strain-strain coupling between each conspecific pair of strains (belonging to the same species) which co-occurred in a community. For each conspecific strain pair, we calculated their temporal abundance trajectories, i.e., their relative abundances at all 8 time points (described above), and then measured the magnitude of the Pearson correlation coefficient between them. We measured only the magnitude of each correlation (regardless of its *P* value) because we were interested only in the extent of covariance between two conspecific strains, and not in the statistical significance of their association. We verified that using a nonparametric covariance measure such as the Spearman correlation did not affect our results (Fig. S9).

#### Interactions between strains and species

We measured putative signatures of interaction in each community separately at the level of species and strains. At each level (say species), we first calculated the temporal abundance trajectories of all members, i.e., their relative abundances at all 8 time points (described above). Then, for each inter-species pair, we then measured the Pearson correlation coefficient between their abundance trajectories (similar results with Spearman correlations, Fig S10). We used the magnitude of the correlation as a proxy for interaction strength. Relative abundance data can exhibit spurious correlations because of finite sampling and the constraint that abundances must sum up to 1. To account for this while detecting the presence of an interaction, we calculated the statistical significance of each correlation against an expected correlation distribution, described as follows.

#### Neutral model to detect interactions

To estimate the expected correlation distribution between the abundance trajectories of two species (or strains) in a community, we used a neutral model to simulate community abundances. The neutral model made two key assumptions: the abundance fluctuations of each member were independent of each other (as expected when there are no interactions) and followed a Gamma distribution, which empirically describe abundance fluctuations in microbial communities (*27, 28*). For each species *i*, we used the observed mean 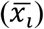 and variance 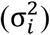 of its abundance, *x*_*i*_, from all our communities to fix the parameters of its expected abundance distribution 𝒫_*i*_(*x*) as follows:

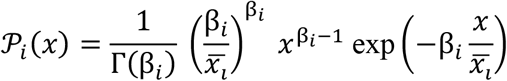

Here, 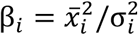. We then simulated the abundance trajectories of each community with the same species or strains as laboratory communities. At each time point, we randomly picked the abundance of each community member from its expected abundance distribution. After picking the abundance of all members, we renormalized them (divided each abundance by the total sum) to obtain relative abundances. We then measured the correlations between each pair of species (or strains) using this synthetic abundance data, using the same procedure described for real data. We repeated the simulations and correlation measurements 1,000 times to build an expected correlation distribution. To measure the statistical significance of a specific correlation observed between two species or strains, we calculated its *P* value using this expected distribution, i.e., the probability of obtaining this correlation by chance. We called an interaction “present” if its *P* value was below 0.05. Since the expected distribution already accounted for multiple comparisons, we did not perform an additional correction.

#### Null model to control for phylogenetic association

To control for the association between species and strains while comparing species-species and strain-strain interaction strengths, we calculated the fraction of cases where strains were more interactive than species in a null model. To do this, we shuffled the association between species and strains by redistributing strains across species. In each community, we randomly shuffled strain labels, thus splitting conspecific strains across different species and coalescing inter-specific strains into the same species. We repeated our measurement of strain-strain and species-species interaction strengths on these shuffled data and calculated the fraction ℱ of cases where the magnitude of correlation between a pair of strains from different (relabeled) species was higher than the correlation between the (relabeled) species themselves. We repeated this shuffling (permutation) 1,000 times, thus obtaining a null distribution 𝒫(ℱ) and a corresponding *P* value for our experimentally measured fraction ℱ_*obs*_ = 76%, i.e., the probability of observing a fraction ℱ equal to or greater than 76% (Fig. S6).

### Code availability

Analysis tools and code are available at: **https://github.com/eltanin4/pitcher_plant_ecoevo**.

### Data availability

Raw sequencing reads are available in the NCBI Sequence Read Archive (BioSample SAMN17005333). Assembled genomes have been deposited in the NCBI GenBank database (BioProject PRJNA682646). Genome metadata and accession numbers are provided in Supplementary Table S1.

## Acknowledgements

We thank Michelle Oraa Ali for their illustration in Fig. 1. This work was supported by the Gordon and Betty Moore Foundation Physics of Living Systems Fellowship grant # GBMF4513 (A.G.), the Human Frontiers Science Program grant # LT000643/2016-L (G.E.L.), the James S. McDonnell Foundation Postdoctoral Fellowship award # 220020477 (L.S.B.), the NSF-DEB grant # 1655983 (O.X.C.), and the Simons Foundation Collaboration: Principles of Microbial Ecosystems (PriME) award # 542395 (O.X.C.).

## Author contributions

A.G.: Conceptualization (data analysis), methodology, software, formal analysis, data curation, writing – original draft, writing – review and editing, visualization. L.S.B.: Conceptualization (experimental design), methodology, data collection, data curation. G.E.L.: Conceptualization (experimental design), methodology, software, data collection, data curation. L.L.: Data collection. O.X.C.: Conceptualization (data analysis), methodology, writing – review and editing, supervision, funding acquisition.

