## Supplementary Material for "Interactions between strains govern the eco-evolutionary dynamics of microbial communities"

### Supplementary Figures

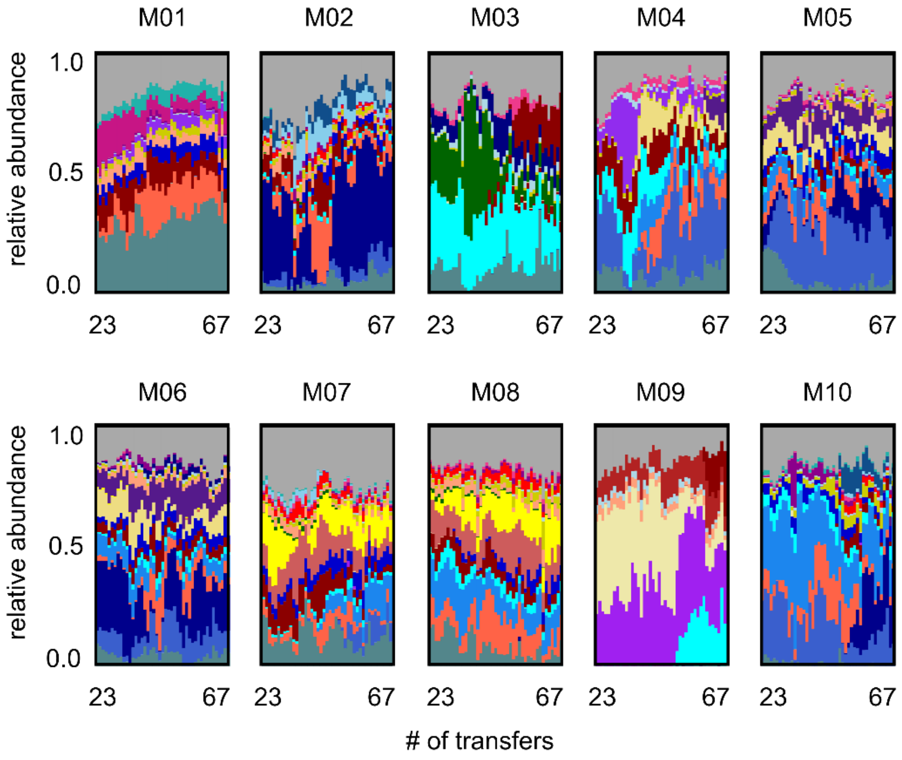


**Figure S1: Long-term species dynamics of all 10 experimental microbial communities.** Stacked bar plots show the composition of all 10 communities (M01 to M10) at the ASV (species) level sampled at each transfer; each color corresponds to a unique ASV which we tracked further using metagenomic sequencing.

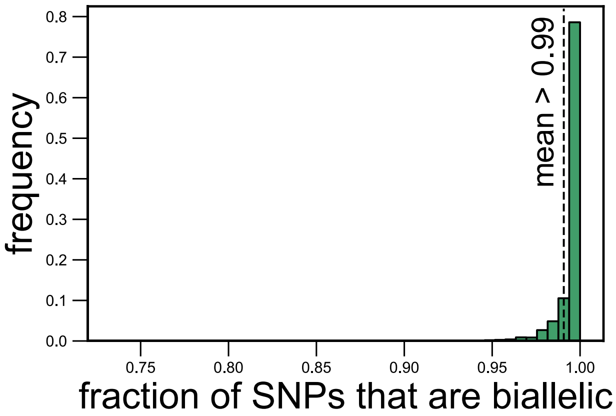


**Figure S2: Most (>99%) SNPs are bi-allelic (have two alleles).** Histogram showing the fraction of SNPs corresponding to each species in each metagenomic sample for which we could detect only two alleles. The dashed line shows the mean fraction of SNPs that are biallelic (0.99).

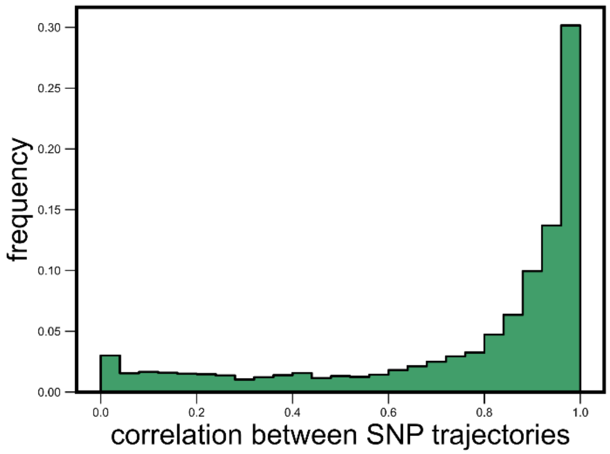


**Figure S3: SNP trajectories within a species are highly correlated.** Histogram showing the distribution of the Pearson correlation coefficient between all pairs of SNP trajectories belonging to the same species in the same community, measured across all species and communities. The mean correlation between SNPs was 0.78.

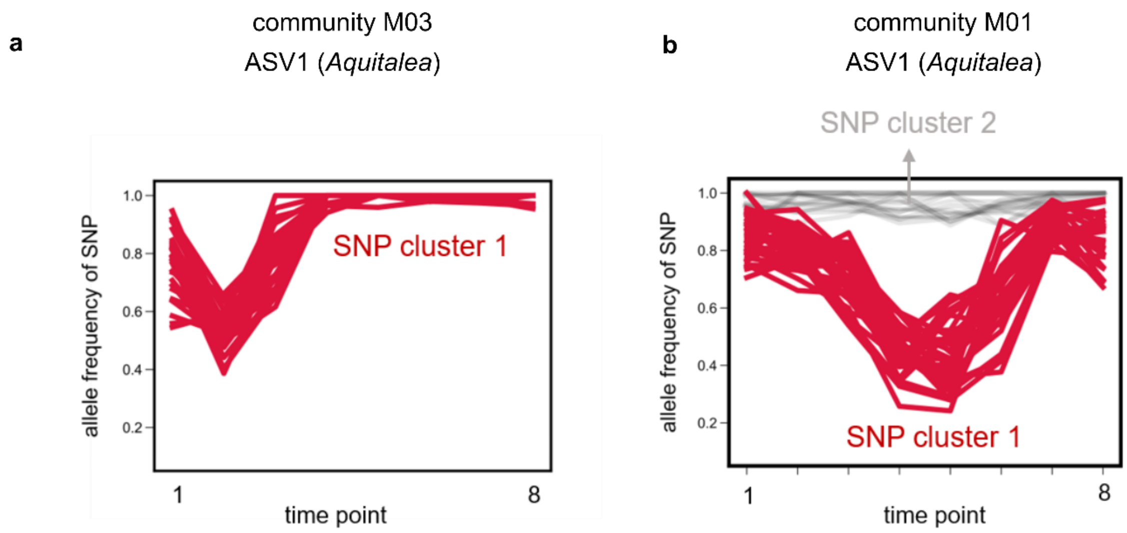


**Figure S4: SNPs within strains tightly cluster together.** Examples of allele frequency trajectories of SNPs belonging to the same species in the same community for ASV1, belonging to the genus *Aquitalea*, in communities **(a)** M03 and **(b)** M01, respectively. SNP trajectories are colored and marked according to the cluster they were identified to be part of using *k*-means clustering (see Methods). (a) shows an example where only one cluster was identified, whereas (b) shows an example of two clusters. SNP cluster 2 consisted of SNPs with an allele frequency near 1, and were therefore assumed to be shared between and common to both strains (but different from the reference genome; see Methods).

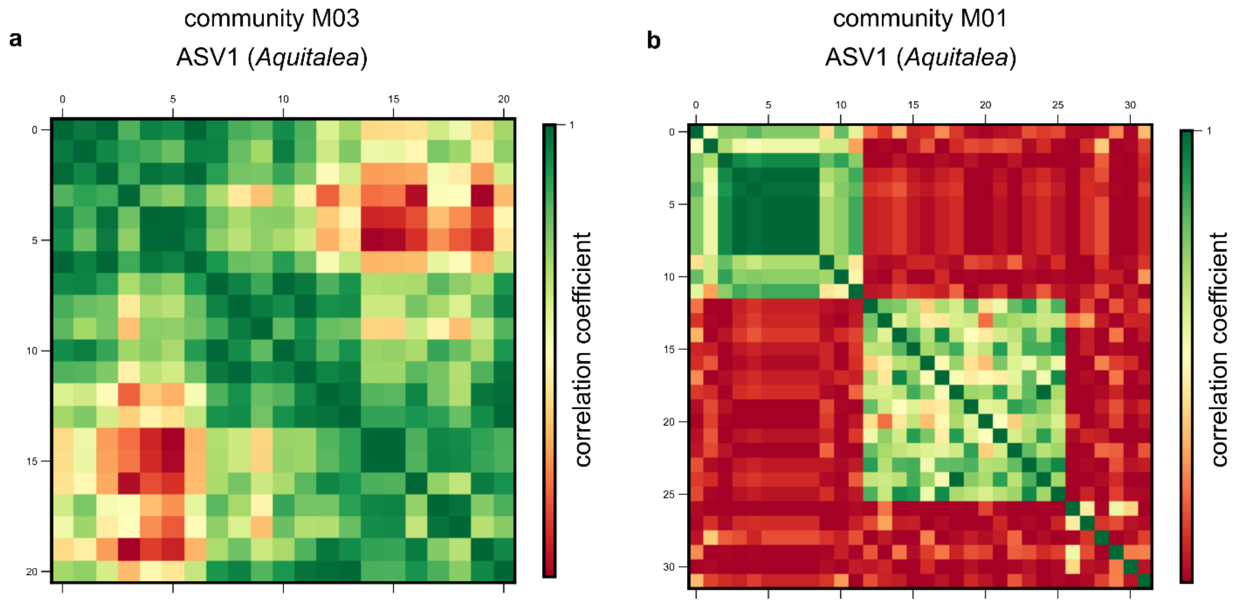


**Figure S5: SNP clusters are robust to alternate clustering methods.** Examples of correlation matrices between SNP trajectories belonging to the same species in the same community for ASV1, belonging to the genus *Aquitalea*, in communities **(a)** M03 and **(b)** M01, respectively. Each row and column represents a distinct SNP trajectory for that species, and colors represent the Pearson correlation coefficient between them. SNPs are ordered by according to clusters identified using a hierarchical clustering algorithm (see Methods).

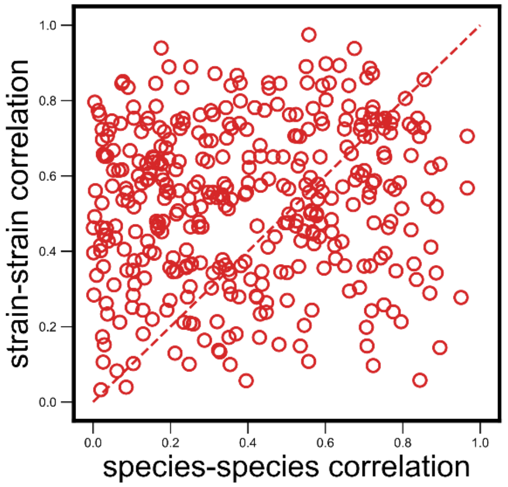


**Figure S6: Null model where we shuffled species-strain associations does not show the observed strain-specificity.** Scatter plot of the dynamical correlation between species in a community and the highest correlation between their corresponding strain pairs (similar to Fig. 3b) but with species and strain associations shuffled within a community (see Methods). Each point represents one species in one of the 10 communities, but is composed of two randomly chosen and unrelated strains from the community. Doing so results in a much lower fraction of correlations that are higher for strains (in this example, 62%, compared with 76% for real data; mean shuffled data fraction was 64%). The probability of observing a fraction greater than or equal to the observed fraction in Fig. 3b by removing any species-strain associations was less than 0.1% ($P < 10^{-3}$).

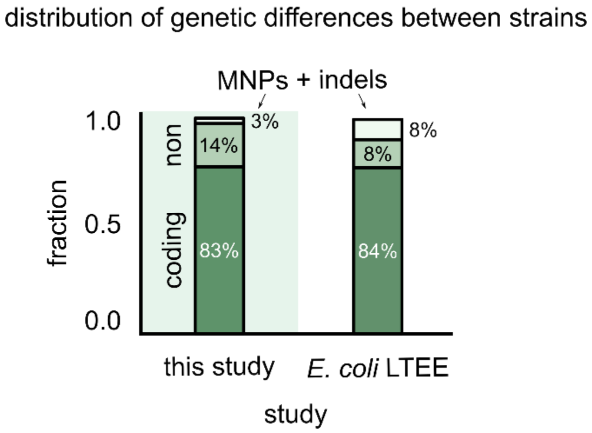


**Figure S7: Most SNPs which differentiate strains are in coding regions.** Stacked bar plots showing the distribution of genetic differences between strains in our communities (left) and variants in the *E. coli* long term evolution experiment after 60,000 generations (LTEE) (data from Good et al (*9*)). The top stack shows all multiple nucleotide polymorphisms (MNPs) and insertions-deletions (indels); the middle stack shows the single nucleotide polymorphisms (SNPs) in noncoding regions of strain genomes, and the bottom stack shows SNPs in coding regions.

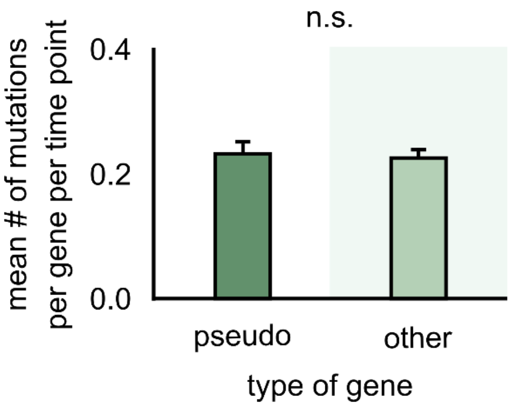


**Figure S8: Mutations accumulate at a similar rate in both pseudogenes and other genes.** Bar plots showing the average number of mutations detected in a pseudogene (left) and any other gene (right) in strains within our 10 communities. The number of mutations was measured per gene per time point to correspond to a rate of mutation accumulation per gene per time point. (Each time point corresponds to about 40 microbial generations.) Error bars show standard error of the mean (s.e.m.). n.s. indicates that there was no significant difference in mutation rate between pseudogenes and other genes ($P > 0.05$, according to a two-sample Student's *t*-test).

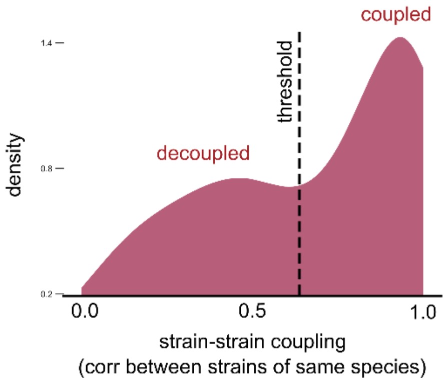


**Figure S9: Strain-strain coupling distribution is robust to using an alternate measure.** Distribution of the strain-strain coupling across all species and communities (similar to Fig. 2b) but measured using the nonparametric Spearman correlation coefficient (see Methods); dashed line shows the internal inflection point of the distribution, separating decoupled and coupled strains.

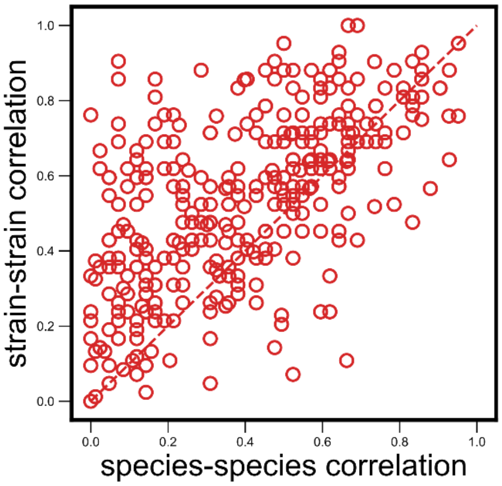


**Figure S10: Strain-specific interactions are stronger even when using an alternate measure.** Scatter plot of the dynamical correlation between species in a community and the highest correlation between their corresponding strain pairs (similar to Fig. 3b) but correlations measured using the nonparametric Spearman correlation coeffcient (see Methods). Each point represents one species in one of the 10 communities. Similar to the fraction of strain-specific interactions observed in Fig. 3b (76%), we see a large fraction of interaction strengths (80%) skewed towards strains.

### Supplementary Tables

**Table S1:** Metadata and accession numbers for all 33 assembled genomes used in the study.

**Table S2:** Set of SNP locations and corresponding gene annotations for members of an example community M04.
